# Complexin suppresses spontaneous exocytosis by capturing the membrane-proximal regions of VAMP2 and SNAP25

**DOI:** 10.1101/849885

**Authors:** J. Malsam, S. Bärfuss, T. Trimbuch, F. Zarebidaki, A.F.-P. Sonnen, K. Wild, A. Scheutzow, I. Sinning, J.A.G. Briggs, C. Rosenmund, T.H. Söllner

## Abstract

The neuronal protein complexin contains multiple domains that exert both clamping and facilitatory functions to tune spontaneous and action potential triggered synaptic release. We address the clamping mechanism and show that the accessory helix of complexin arrests the assembly of the soluble N-ethylmaleimide-sensitive factor attachment protein receptor (SNARE) complex that forms the core machinery of intracellular membrane fusion. In a reconstituted fusion assay, site- and stage-specific photo-cross-linking reveals that prior to fusion the complexin accessory helix laterally binds the membrane-proximal C-terminal ends of SNAP25 and VAMP2. Corresponding complexin interface mutants selectively increase spontaneous release of neurotransmitter in living neurons, implying that the accessory helix suppresses final zippering/assembly of the SNARE four-helix bundle by restraining VAMP2 and SNAP25.

## Introduction

Signal propagation between neurons relies on fast quantal release of neurotransmitters from the presynaptic terminal into the synaptic cleft. An incoming action potential elicits the influx of Ca^2+^ into the nerve terminal, which instantly triggers the fusion of neurotransmitter-filled synaptic vesicles, docked at the active zone of the presynaptic plasma membrane (Rothman, 2014; Sudhof, 2013). The underlying core fusion machinery is formed by trans v-/t-SNARE complexes (SNAREpins) bridging the vesicular and plasma membrane (Söllner et al., 1993; Weber et al., 1998). In a zipper-like manner, 16 layers of the α-helical SNARE motif assemble into a four-helix bundle pulling the opposing membranes together to drive bilayer fusion (Figure 1A) (Stein et al., 2009; Sutton et al., 1998). The v-SNARE VAMP2/synaptobrevin, anchored to synaptic vesicles, provides one helix, and the t-SNARE (syntaxin1 and SNAP25), localized to the pre-synaptic plasma membrane, contributes three helices. SNAREpin formation is precisely controlled by regulatory proteins accelerating and arresting distinct assembly steps (Brunger et al., 2018; Jahn and Fasshauer, 2012; Sudhof, 2013). Crucially, the Ca^2+^ sensor synaptotagmin1 (Syt1), which is anchored to synaptic vesicles and the small cytosolic protein complexin (Cpx) arrest SNAREpin assembly at a late metastable state, a prerequisite for the hallmark fast Ca^2+^-synchronization (Fernandez-Chacon et al., 2001; Geppert et al., 1994; McMahon et al., 1995; Melia, 2007; Mohrmann et al., 2015; Trimbuch and Rosenmund, 2016). In the absence of Syt1, spontaneous release events are increased and evoked exocytosis is impaired (DiAntonio and Schwarz, 1994; Geppert et al., 1994; Littleton et al., 1994; Pang et al., 2006). Syt1 contains two cytosolic C2 domains (C2A, C2B), which bind Ca^2+^, anionic phospholipids, and SNAREs, likely forming oligomeric assemblies and restraining SNAREpins to arrest the prefusion stage in the absence of Ca^2+^ (Bello et al., 2018; Brose et al., 1992; Davletov and Sudhof, 1993; de Wit et al., 2009; Li et al., 2019; Perin et al., 1990; Zhou et al., 2017). In the presence of Ca^2+^, the C2 domains deform the membrane, exerting force on the SNAREpins and membrane fusion is triggered (Chapman and Davis, 1998; Fernandez et al., 2001; Fernandez-Chacon et al., 2001; Hui et al., 2009; Martens et al., 2007; Wang et al., 2016).

**Figure 1.**
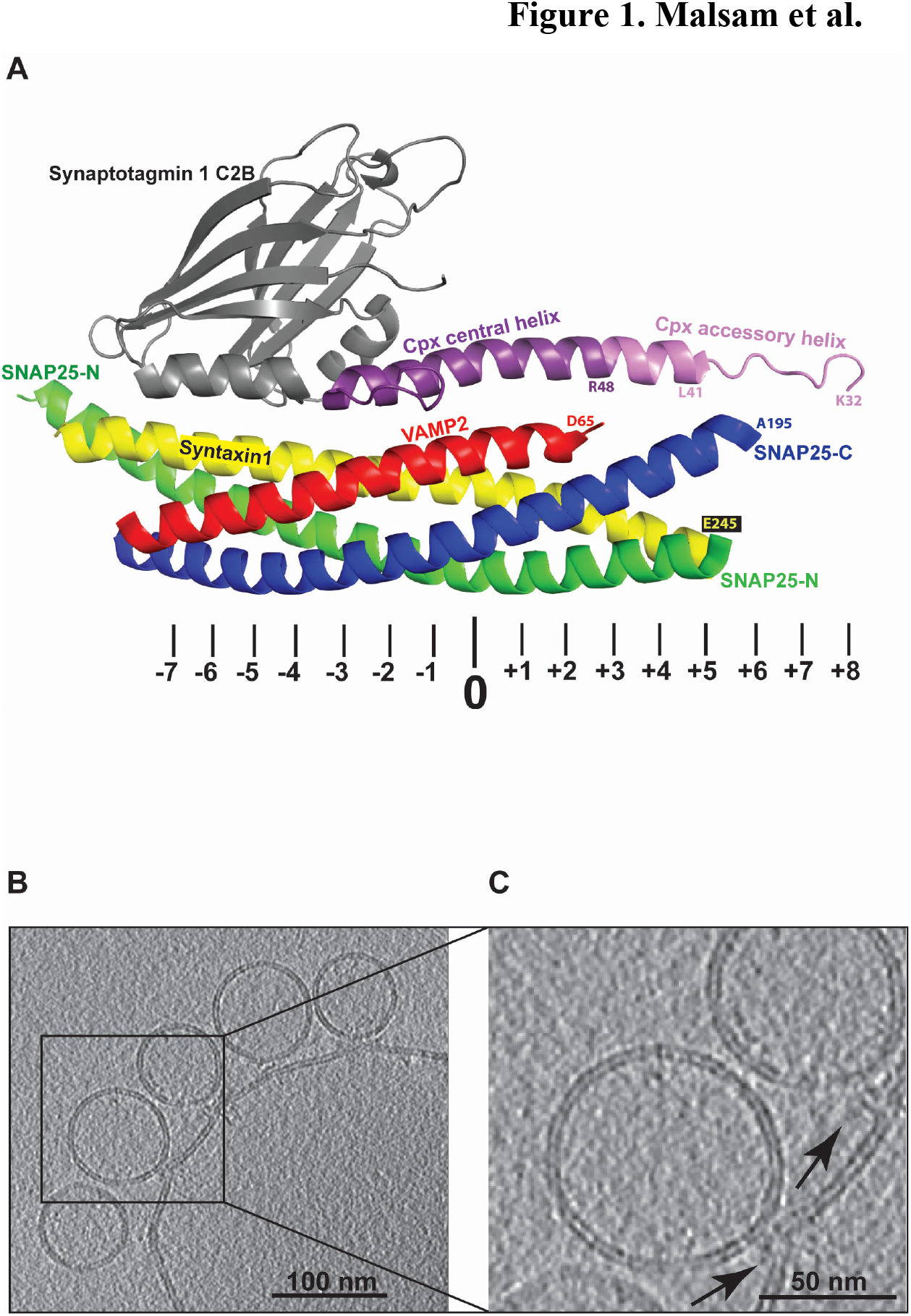
Cpx-Syt1 primed SNAREpins and docked vesicles. (A) Crystal structure of the membrane-distal Cpx-Syt1-SNARE interface (modified from (Zhou et al., 2017)). The positions of the hydrophobic layers and the central ionic 0-layer within the SNARE motifs are indicated below. The second C2B binding site facing SNAP25 is not shown. (B) Slice through a cryo-tomogram of docked SUVs, whose fusion can be triggered by Ca^2+^. GUVs containing full-length t-SNARE complexes (syntaxin1/SNAP25) were mixed with SUVs containing Syt1 and the v-SNARE VAMP2 in the presence of Cpx wt. Samples were preincubated for 30 minutes on ice to efficiently accumulate the Cpx-stabilized prefusion intermediate. (C) Magnification of area outlined in B. Arrows indicate the position of putative SNAREpins and associated proteins.

Like Syt1, complexin controls spontaneous and evoked exocytosis. To which degree complexin suppresses/stimulates spontaneous release events is controversial because knock-out or knock-down manipulations yield diverse results and show neuron-specific differences (Yang et al., 2013). In addition, in vertebrates, such as *Mus musculus*, the stimulatory role seems to dominate while in invertebrates, such as *Drosophila melanogaster* and *Caenorhabditis elegans*, the inhibitory function dominates (Hobson et al., 2011; Huntwork and Littleton, 2007; Lin et al., 2013; Lopez-Murcia et al., 2019; Maximov et al., 2009; Pang et al., 2006; Reim et al., 2001; Strenzke et al., 2009; Wragg et al., 2017; Xue et al., 2007). Complexin consists of an N-terminal stimulatory region (aa 1-25) that activates Ca^2+^-triggered release, an inhibitory accessory helix (aa 26-47) that suppresses spontaneous release, a central SNARE-binding helix (aa 48-74), and a largely unstructured C-terminal region (aa 75-134) (Buhl et al., 2013; Cho et al., 2014; Cho et al., 2010; Gong et al., 2016; Iyer et al., 2013; Kaeser-Woo et al., 2012; Kummel et al., 2011; Lai et al., 2016; Lai et al., 2014; Martin et al., 2011; McMahon et al., 1995; Snead et al., 2014; Xue et al., 2010; Xue et al., 2007; Yang et al., 2015). The C-terminal region contains a short amphipathic helix which binds high curvature membranes and modulates the species-specific inhibition (Gong et al., 2016; Seiler et al., 2009; Snead et al., 2014; Wragg et al., 2017; Wragg et al., 2013; Zdanowicz et al., 2017). Interestingly, a recent study using magnetic tweezers and truncated SNARE proteins revealed that amino acids 1-31 of CpxI slow down the zippering of the linker regions connecting the SNARE motifs to their transmembrane regions (Shon et al., 2018). Thus, the exact nature and the molecular interactions of the membrane proximal, N-terminal 47 amino acids, containing both activating and suppression functions, still remain unclear.

Crystal and NMR structures of SNARE-Syt1, SNARE-Cpx, and SNARE-Cpx-Syt1 complexes have provided critical information about intermolecular interactions (Bracher et al., 2002; Brewer et al., 2015; Chen et al., 2002; Zhou et al., 2015; Zhou et al., 2017). However, many of these structures reflect the postfusion stage or are based on truncated proteins to mimic potential prefusion stages. A recent crystal structure of a truncated, partially assembled SNARE complex revealed a Syt1-Cpx-SNARE interface localized around the membrane-distal region of the SNAREpin (Zhou et al., 2017). A second Syt1 C2B domain simultaneously binds the other side of the SNAREpin at the central region. Both interactions are required for synchronized evoked exocytosis. In contrast, the organization of the membrane-proximal SNARE regions and, in particular, the biophysical interactions of the inhibitory accessory helix in the prefusion stage, remain elusive and are still highly debated. Several potential models for the action of Cpx have been proposed. These include electrostatic repulsion of negatively charged membranes, stabilization of the central helix by the accessory helix, and direct Cpx-SNARE interactions, also linking SNARE complexes (Choi et al., 2016; Giraudo et al., 2009; Krishnakumar et al., 2015; Lu et al., 2010; Prinslow et al., 2017; Radoff et al., 2014; Trimbuch et al., 2014; Xue et al., 2010; Zdanowicz et al., 2017).

## Results

### Identification of interacting partners of the CpxII accessory helix by site- and stage-specific photo-cross-linking

Since Ca^2+^ triggers membrane fusion on a sub-millisecond time scale, we reasoned that only a few molecular rearrangements may suffice to release *a priori* weak constraints, which the Cpx accessory helix imposes on its putative binding partners. To covalently trap the complexin-SNARE interactions in the metastable prefusion conformation, we used a cross-linking approach to probe the local environment of the accessory helix on a single amino acid level with a space resolution of about 3 Å, both in the pre- and postfusion state. Among the four mammalian complexin paralogs, we chose to analyze CpxII, which controls exocytosis in the brain and in other tissues (Reim et al., 2001; Yamada et al., 1999). We took advantage of a well-characterized reconstituted proteoliposome fusion assay that allows the accumulation of prefusion Cpx-clamped SNAREpins and that upon Ca^2+^-addition triggers fusion, culminating in fully assembled postfusion cis-SNARE complexes (Bharat et al., 2014; Malsam et al., 2012). Each of the 22 amino acids of the CpxII accessory helix (aa 26-47) was exchanged for the photo-activatable, unnatural amino acid benzoyl-phenylalanine (BPA)(Young et al., 2010). In the absence of photo-activated crosslinking, like in a standard mutagenesis approach, the functional contribution of each amino acid can be tested. Upon photoactivation, interaction partners can be identified and SNAREpin zippering may become irreversibly arrested. Prefusion SNAREpins were accumulated by incubating small VAMP2/Syt1-containing liposomes (SUVs) mimicking synaptic vesicles with large syntaxin1/SNAP25-containing liposomes (GUVs) mimicking the pre-synaptic plasma membrane in the presence of CpxII wt or the respective single BPA mutations. Figure 1B shows a slice through a cryo-tomogram of this Cpx-stabilized prefusion intermediate accumulating docked v-SNARE/Syt1 SUVs on the surface of a t-SNARE GUV. Note the electron-dense regions between the two opposing membranes at high magnification (arrows), likely representing arrested SNAREpins and associated proteins (Figure 1C). At some docking sites, the GUV surface protrudes towards the SUV, suggestive of a constrained readily releasable state (Bharat et al., 2014).

To probe the local environment of the Cpx-accessory helix within the prefusion SNAREpin, we subjected samples to UV (365 nm)-irradiation, followed by Western blotting and immuno staining. The detection of cross-linked products with an anti-Cpx antibody revealed specific cross-link bands at 29, 32, 41 kDa and a weaker band at 61 kDa (Figure 2A). The 32 kDa band indicates the presence of Cpx dimers formed by cross-links involving in particular the N-terminus of the accessory helix, which may point towards the site of fusion pore formation. Specific products detected with the anti-Cpx antibody were co-recognized by anti-VAMP2, anti-SNAP25 and anti-Syt1 antibodies, respectively (Figure 2A), identifying inter-subunit cross-links. No Cpx cross-link products could be detected using a syntaxin1-specific antibody. Interestingly, the formation of VAMP2- and SNAP25-Cpx cross-links follows a defined periodicity: pairs of two adjacent Cpx residues are linked to either VAMP2 or SNAP25 in an alternating pattern, indicating the localization of these SNAREs along two opposite sides of the Cpx accessory helix. Indeed, viewed along the helix axis, the crosslinks show that the Cpx accessory helix is sandwiched between SNAP25 and VAMP2 (Figure 2A). In summary, the data reveals the presence of a bipartite Cpx interface prior to fusion, which follows a helical pattern and captures the C-termini of SNAP25 and VAMP2, likely preventing SNARE complex zippering.

**Figure 2.**
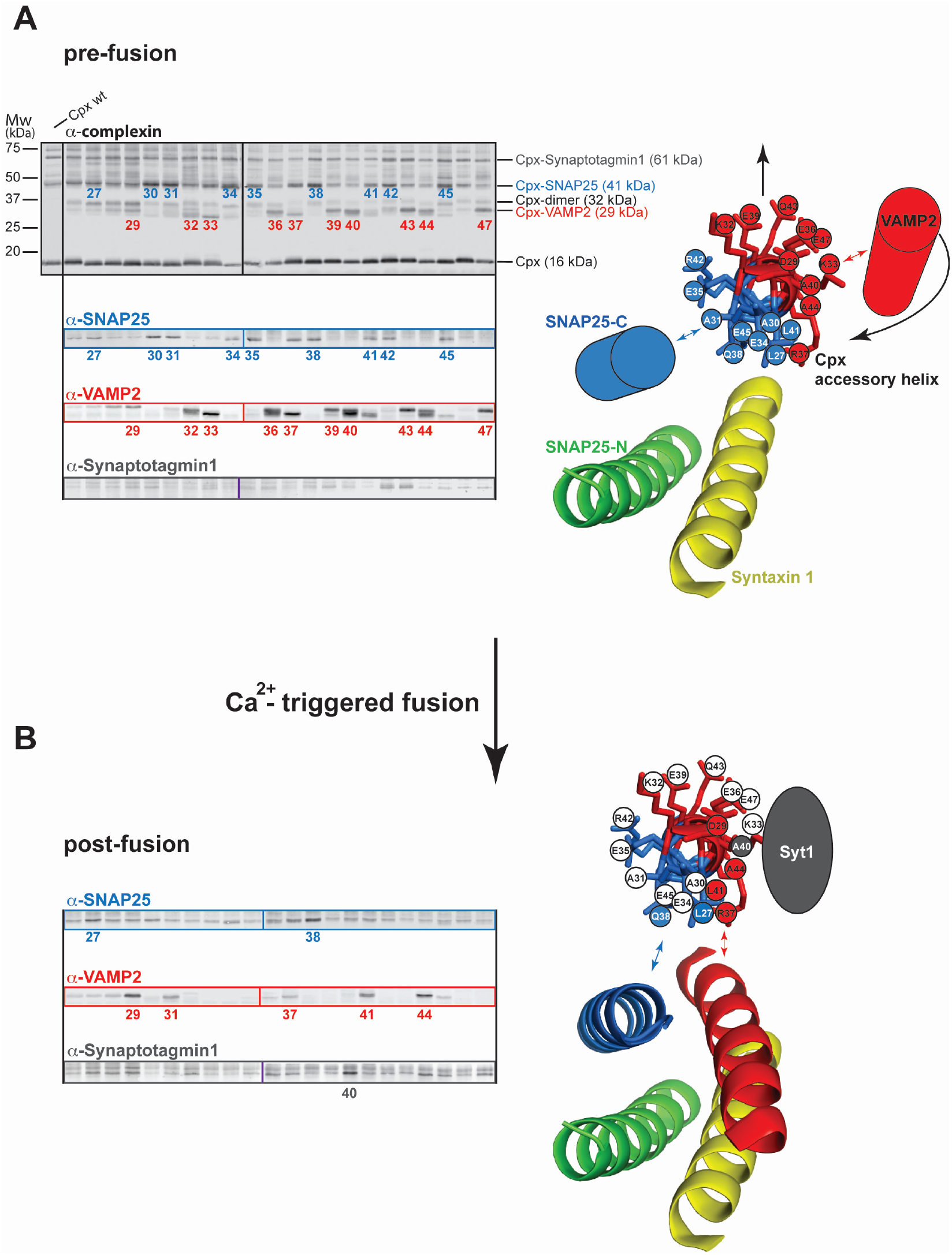
Mapping interactions of the CpxII accessory helix with the fusion machinery by site- and stage-specific cross-linking. GUVs containing the full-length t-SNARE complex (syntaxin1/SNAP25) were mixed with SUVs containing Syt1 and the v-SNARE VAMP2 in the presence of CpxII wt or CpxII-BPA mutants. Samples were preincubated for 30 minutes on ice to accumulate docked SUVs linked to the GUVs by Syt1 and trans-SNARE complexes. UV irradiation of the reaction mix was performed before (prefusion) and after triggering fusion (postfusion) with 100 μM calcium at 365 nm for 15 seconds on ice. Cross-link products were analyzed by Western blotting using the indicated antibodies. Positions of identified cross-link products are indicated by the molecular identity. (A) Cross-link products identified at the prefusion stage (left panels). View along the axis of the CpxII accessory helix shows interactions with SNAP25 (blue) and VAMP2 (red) (right panel). The membrane-proximal regions of VAMP2 and SNAP25 are depicted as cylinders, because their prefusion structures are not known, but may form α helical structures. (B) Cross-link products identified at the postfusion stage and structural model. See also Figure S1

If this pattern is specific for the prefusion stage it should change postfusion. Thus, the systematic cross-link approach was repeated with samples, where liposome fusion had been triggered with 100 μM free Ca^2+^ prior to UV treatment (Figure 2B). The Cpx cross-link pattern changes profoundly. Postfusion, cross-link formation is restricted to a narrow region of the Cpx helical wheel that projects towards different VAMP2 and SNAP25 interfaces, which is consistent with the position of Cpx in the postfusion crystal structure (Figure 2B) (Chen et al., 2002). No obvious cross-links to Syt1 were observed in the prefusion state. However, at the postfusion stage, a single prominent cross-linked product of 61 kDa is generated by the Cpx residue A40-BPA, which was recognized by both an anti-Syt1 - and an anti-Cpx antibody (Figure 2B and Figure S1). Since Cpx A40-BPA displays a prominent cross-link with VAMP2 at the prefusion state (Figure 2A) this finding suggests that a fusion/Ca^2+^-dependent conformational rearrangement now allows Syt1 to interact with Cpx at its VAMP2 prefusion interface.

To narrow down the region of SNAP25 that cross-links to the Cpx accessory helix in the prefusion state, we repeated the above experiments using C-terminally truncated SNAP25 constructs terminating either at layer +6 (SNAP25 amino acids1-194) or at layer +8 (SNAP25 amino acids 1-200) of the second SNARE motif (Figure S2A). When using SNAP25 1-194 we saw no SNAP25-positive Cpx cross-link products, localizing the Cpx-interacting region to the 12 C-terminal residues of SNAP25 (Figure S2B). To map the VAMP2 interacting region, we performed a proteolytic analysis of VAMP2 cross-linked to CpxR37BPA (Figure S2C). UV-irradiated samples were subjected to site specific proteolytic cleavage by botulinum toxins B and D, cleaving VAMP2 at distinct sites. The proteolytic pattern allocates the cross-linked site to the last three layers (+6, +7 and +8) of the VAMP-2 SNARE motif. Thus, the binding sites of the Cpx accessory helix can be confined to the membrane-proximal C-terminal ends of SNAP25 and VAMP2. Interestingly, single molecule optical tweezer measurements revealed that in isolated t-SNARE complexes layers +5 to +8 are frayed and thus may become a natural target for Cpx accessory helix binding (Zhang et al., 2016). In turn, large parts of the Cpx accessory helix seem to be unstructured in the absence of the C-terminal SNARE regions (see also Figure 1A) (Maximov et al., 2009; Trimbuch et al., 2014; Zhou et al., 2017). Therefore, the cross-linking data in combination with previous work suggest that the Cpx accessory helix and the membrane proximal ends of SNAP25 and VAMP2 mutually influence each other’s structure, likely forming a short three-helix bundle, which opposes the final zippering of the four-helix SNARE bundle (Figure 2). Such a model would be overall consistent with the observation that Cpx affects the SNARE assembly state and that an extended helical structure of Cpx is required for fusion arrest (Choi et al., 2016; Yin et al., 2016).

### Site specific arrest of the fusion machinery by cross-linking of the CpxII accessory helix inhibits Ca^2+^-triggered membrane fusion in a reconstituted assay

In functional terms, the site-specific cross-links should fix the complex in the prefusion state substantially interfering with subsequent Ca^2+^-triggered fusion events. We therefore compared untreated and UV-exposed samples in a lipid-mixing assay to monitor the effect of cross-link formation on liposome fusion kinetics. Irradiation of a control reaction containing CpxII wt confirmed that nonspecific effects of UV exposure are negligible. In contrast, UV cross-linking of CpxII BPA mutants results in a variety of phenotypes showing moderate (A30BPA) or strong inhibition of fusion kinetics (A40BPA) (red curves in Figure 3A). Interestingly, in the absence of UV irradiation, replacement of single native amino-acids by the bulky BPA at a few selective positions in the accessory helix reduced complexin’s clamping function, but did not affect final Ca^2+^-triggered fusion (Figure S3). As a control we introduced BPA at position R48, a residue in the central helix, essential for the overall binding of Cpx to the SNAREpin (Xue et al., 2007). This mutation resulted in a loss of the complexin function, abolishing both the clamp and Ca^2+^-dependent stimulation (Figure 3A).

**Figure 3.**
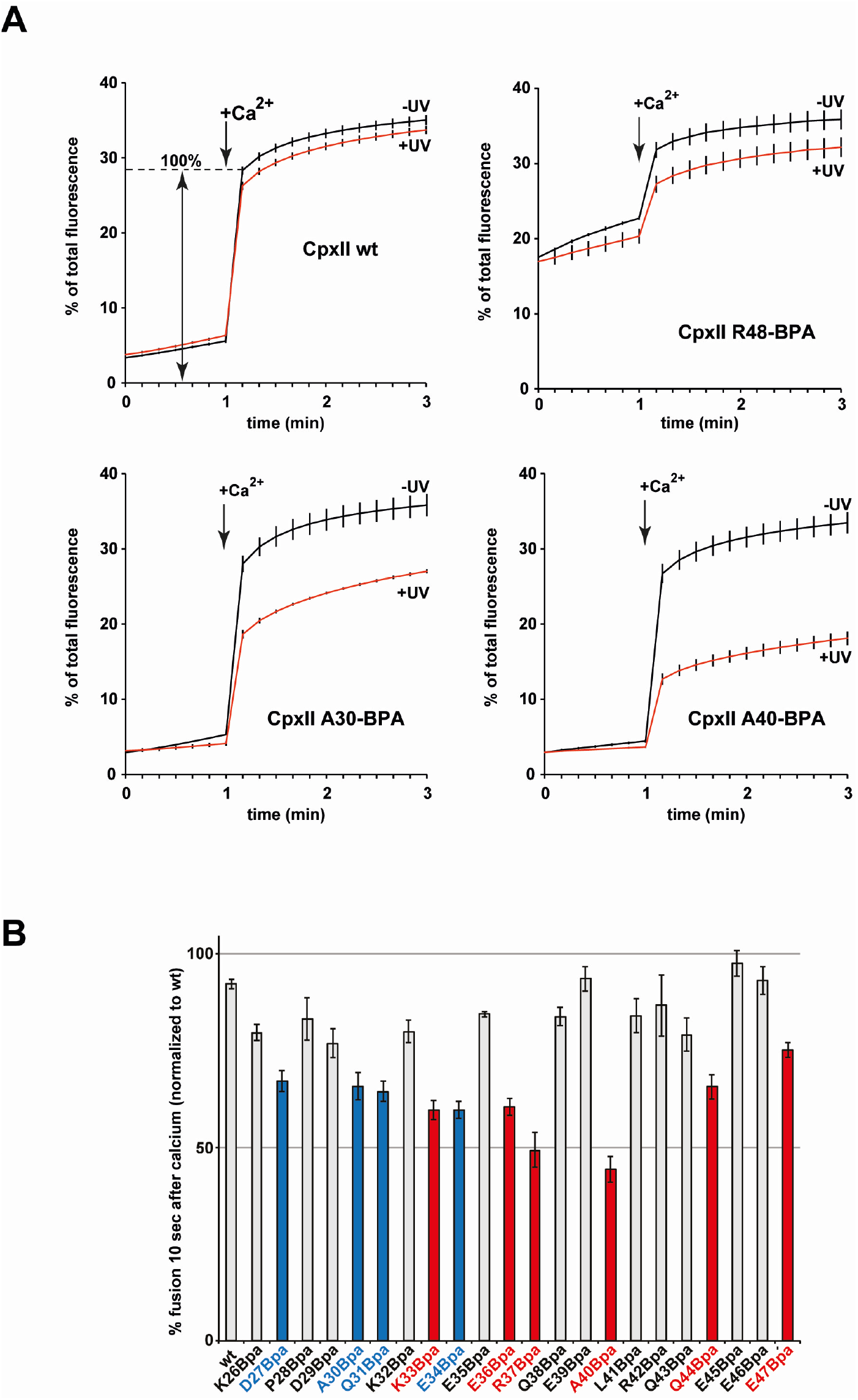
Site specific arrest of the fusion machinery by scanning BPA cross-linking of the CpxII accessory helix. Syntaxin1/SNAP25-GUVs were mixed with Syt/VAMP2-SUVs in the presence of CpxII wt or the indicated Cpx BPA mutants, and incubated for 30 minutes on ice to dock vesicles. The reaction mixes were irradiated at 365 nm for 15 seconds on ice (control reactions without UV irradiation). Subsequently, fusion kinetics were recorded at 37°C for one minute in the absence of Ca^2+^ and the measurement was continued for another two minutes after injection of 100 μM free Ca^2+^ to trigger fusion. (A) Lipid-mixing kinetics of CpxII wt and distinct CpxII BPA mutants +/− UV-irradiation. Please note that CpxII R48BPA impairs SNARE complex binding, resulting in a loss of the clamp and in an elevated starting signal. (B) BPA cross-link scan of the complete accessory helix and the effect on Ca^2+^-triggered fusion (signal change 10s after Ca^2+^ trigger as illustrated in (A) wt). Blue and red bar graphs indicate crosslinks to SNAP25 and VAMP2, respectively (n = 3). See also Figures S3 and S4

We extended the analysis of UV-induced fusion arrest to all remaining residues of the CpxII accessory helix and normalized the data to the maximum fluorescence value of a control reaction containing CpxII wt monitored 10 seconds after the addition of calcium (Figure 3B). As expected, the pattern of UV-induced fusion inhibition correlated to a large extent with those residues of the CpxII accessory helix that formed UV-dependent cross-links with either SNAP25 or VAMP2 with maximum inhibition localized to the two adjacent residues R37 and A40 (compare Figures 3B and 2A).

In addition to the biochemical assay, we analyzed the effect of cross-linking on Ca^2+^ triggered fusion in a morphological assay, using cryo-EM and the CpxA40BPA mutant. In the absence of Ca^2+^, approximately ten SUVs were docked per GUV independent of crosslinking (Figures S4A and S4B). Ca^2+^-addition triggered efficient membrane fusion and approximately one to two SUVs remained docked to the GUV surface. In contrast, UV-irradiation profoundly abrogated Ca^2+^-triggered fusion of the BPA mutant and approximately ten SUVs remained docked per GUV. Thus, crosslinking efficiently arrests membrane fusion. Furthermore, it appears that our rather conservative evaluation of the biochemical data underestimates the inhibitory effects.

### Mutants affecting the SNAP25- and VAMP2-/Cpx interface selectively increase Ca^2+^-independent fusion of proteoliposomes and spontaneous but not evoked neurotransmitter release in neurons

The question arises to which degree the interfaces identified by photo-crosslinking selectively affect fusion clamping. Therefore, we generated Cpx quadruple mutants containing reverse charge exchanges in either SNAP25- or VAMP2-interacting residues and analyzed their effect in the liposome fusion assay (Figure 4A). Fusion reactions containing either the SNAP25- or VAMP2-binding quadruple mutant revealed significant defects in suppressing calcium-independent fusion when compared to Cpx wt (Figure 4A, initial two minutes). Importantly, the mutations did not affect the Ca^2+^-triggered total fusion. The SNAP25-binding mutant even showed slightly increased fusion. Altogether, the data reveal that these quadruple mutants primarily affect fusion clamping.

**Figure 4.**
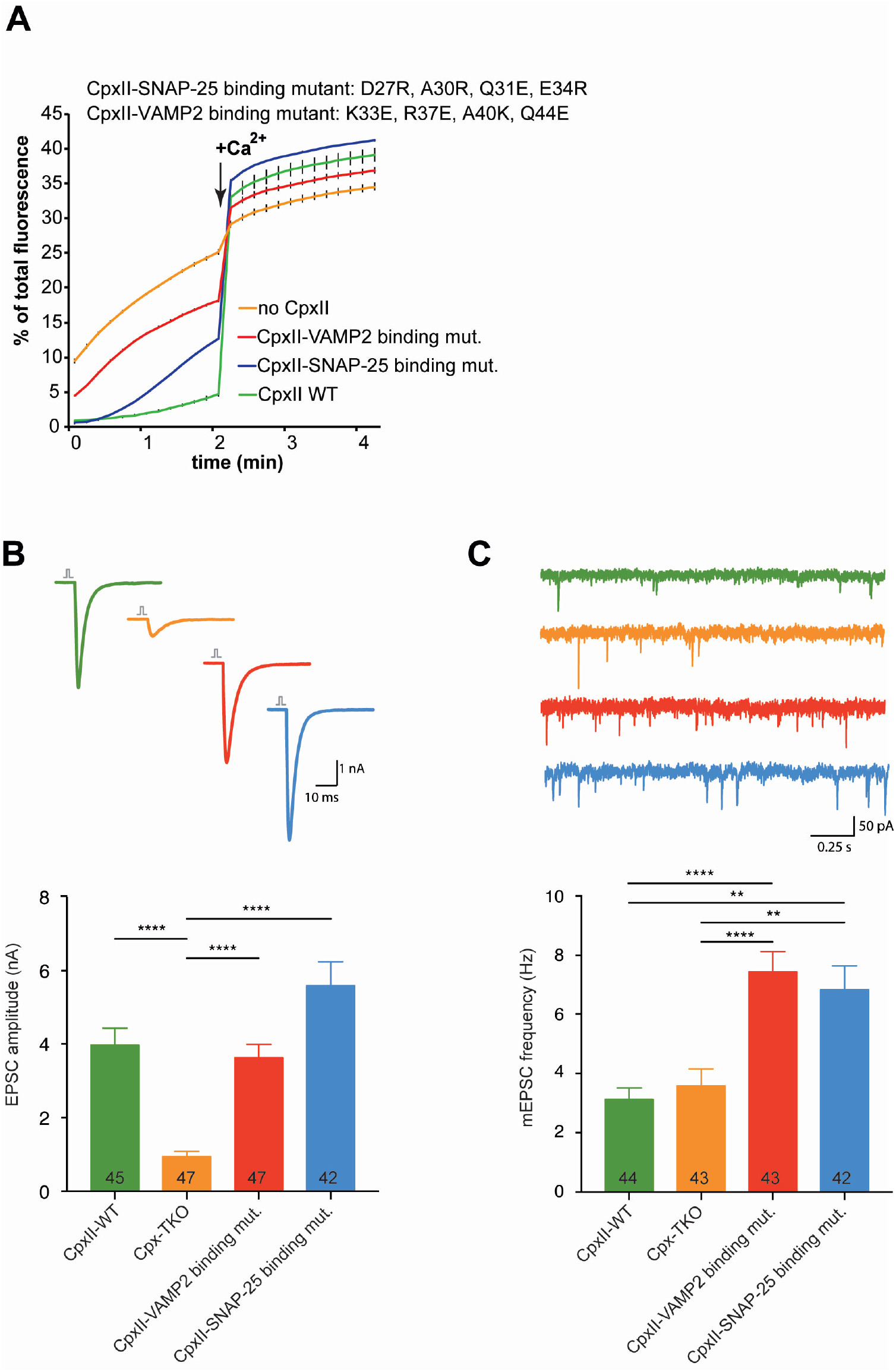
CpxII quadruple mutants of the SNAP25- and VAMP2-binding regions selectively increase Ca^2+^-independent fusion of proteoliposomes and spontaneous but not evoked neurotransmitter release in neurons. (A) t-SNARE-GUVs were mixed with Syt1/VAMP2-SUVs in the absence or presence of either CpxII wt or the indicated mutants. Samples were pre-incubated for 5 minutes on ice and subsequently, fusion kinetics were recorded at 37°C for 2 minutes in the absence of Ca^2+^. Fusion was triggered by injection of 100 μM free Ca^2+^. (B) Cpx I-III KO glutaminergic neurons were transduced with lentivirus containing CpxII wt or the mutants. Analysis of evoked responses by mean EPSC amplitudes. (C) Spontaneous release activity as determined by mean mEPSC frequency. Data information: Error bars indicate s.e.m ** p<0.01, *** p<0.001, **** p<0.0001. See also Figure S5

If the *in vitro* results are physiologically relevant, then in neuronal exocytosis, the same Cpx mutants should predominantly affect spontaneous neurotransmitter release leaving evoked release largely untouched. Therefore, we used lentiviral transduction of wild-type or mutant Cpx to perform rescue experiments on hippocampal glutamatergic neurons from Cpx I-III triple knock-out (TKO) mice (Trimbuch et al., 2014; Xue et al., 2010; Xue et al., 2007; Xue et al., 2008). Western blot analyses showed similar expression levels of all Cpx constructs (Figure S5A). As previously published, the electrophysiological measurements using autapses confirmed that the Cpx TKO profoundly reduced evoked neurotransmitter release, which could be rescued by Cpx wt expression (Figure 4B). Both Cpx quadruple mutants rescued evoked release similar to Cpx wt. Neither Cpx mutants affected the readily releasable pool (RRP) or the release probability (Pvr) (Figures S5B and S5C). Short-term plasticity characteristics, analyzed by a train of action potentials at 50 Hz and 10 Hz, were rescued through expression of the Cpx mutants in the TKO. In line with the rescue of release probability, facilitation in the TKO was converted back to synaptic depression in all mutants tested (Figure S5E and S5F). As established previously, loss of Cpx in hippocampal primary neurons does not increase the rate of spontaneous release (measured as the mini-EPSC (mEPSC) frequency), because strong stimulatory functions of Cpx on release can effectively counter the inhibitory functions (Figure 4C) (Xue et al., 2009; Xue et al., 2008). In contrast, the introduction of either the VAMP2- or the SNAP25-binding mutants into the Cpx TKO neurons increased the spontaneous release by a factor of two (Figure 4C). Thus, the interactions of the accessory helix with both SNARE interfaces specifically govern fusion clamping. Control experiments analyzing the amplitude of single spontaneous release events demonstrate that the Cpx mutants did neither cause alterations in the vesicle size nor neurotransmitter loading, excluding indirect effects on spontaneous exocytosis (Figure S5D).

## Discussion

Overall, our systematic, unbiased approach using site-specific photo-crosslinking revealed that in the prefusion state, the accessory helix of CpxII forms a dual interface with the membrane proximal regions of VAMP2 and the second SNARE motif of SNAP25. To identify these interactions, it was crucial to employ reconstituted full-length proteins in a proteoliposome fusion assay, maintaining the natural membrane constraints, which are characteristic for the prefusion site and thereby also confine the structural organization of the fusion machinery. Our functional studies *in vitro* and in living neurons strongly suggest that both accessory helix interfaces are required for the clamping reaction.

While there is general agreement that complexins stimulate evoked exocytosis, the physiological relevance of suppression of spontaneous fusion in mouse neurons is still debated. The techniques employed, such as neuronal cultures (autapses or interneuronal networks), and the complexin inactivation procedure (knockout or knockdown), may affect the outcome of the measurements (Maximov et al., 2009; Yang et al., 2013). In a recent publication, this issue was re-investigated using acute complexin I depletion in neuronal cultures derived from CpxII/III double knockout mice, which are viable and fertile (Lopez-Murcia et al., 2019; Xue et al., 2008). This conditional depletion largely avoids indirect effects such as collateral perturbations, compensatory processes, or aberrant synaptogenesis. Lopez-Murcia and colleagues showed that the complexin-knockout reduces both spontaneous and evoked release in mouse hippocampal neurons, consistent with our electrophysiological studies using neurons derived from CpxI/II/III triple knockout mice. Strikingly, our rescue experiments using Cpx mutants impaired in binding VAMP2 and SNAP25 reveal that in addition to this stimulatory function of Cpx, the accessory helix mediates a prominent, suppressive clamping function of. For spontaneous release to occur at wild type frequency, synaptic vesicles need to enter a fully primed state and this depends on the activating role of complexins. Our results suggest that the accessory helix acts as a built-in antagonist to restrict this strong activating function, establishing a decisive metastable state. To which degree other domains of complexin, like the C-terminal regions, which bind high curvature membranes and provide clamping functions at least in invertebrates, mechanistically modulate the accessory helix function or act in an independent manner remains to be resolved (Wragg et al., 2017; Xue et al., 2009).

Our study now suggests a simple model for the accessory helix in which the membrane proximal ends of VAMP2 and SNAP25 can form two alternative complexes: i) a three helical bundle with the Cpx accessory helix (Figure 5) or ii) a four-helix bundle with its syntaxin1 and SNAP25 partners allowing further SNARE zippering and initiating membrane fusion. The dual and apparently weak accessory helix-SNARE interactions have important implications: i) by avidity a sufficiently strong clamp is achieved, ii) the weak interactions will likely allow fast release upon arrival of the trigger, and iii) the three interaction partners may influence each other’s structure, thus, generating a reaction intermediate, which may accelerate final SNAREpin zippering when unleashed by the Ca^2+^-Syt1 trigger. The Cpx accessory helix/VAMP2/SNAP25 interaction may also help to position the N-terminus of Cpx relative to the membrane and SNAREpin thereby controlling its stimulatory role. How the Ca^2+^ influx triggers the structural transition remains to be determined. Direct Cpx-Syt1 interactions, as revealed by the crosslink in position 40 of CpxII, may play a role. Whatever the exact release mechanism may be, the identified Cpx-SNARE interactions make profound contributions to clamp spontaneous neurotransmitter release in the central nervous system, likely setting the correct threshold for neurotransmission and for proper signal transduction in the brain.

**Figure 5.**
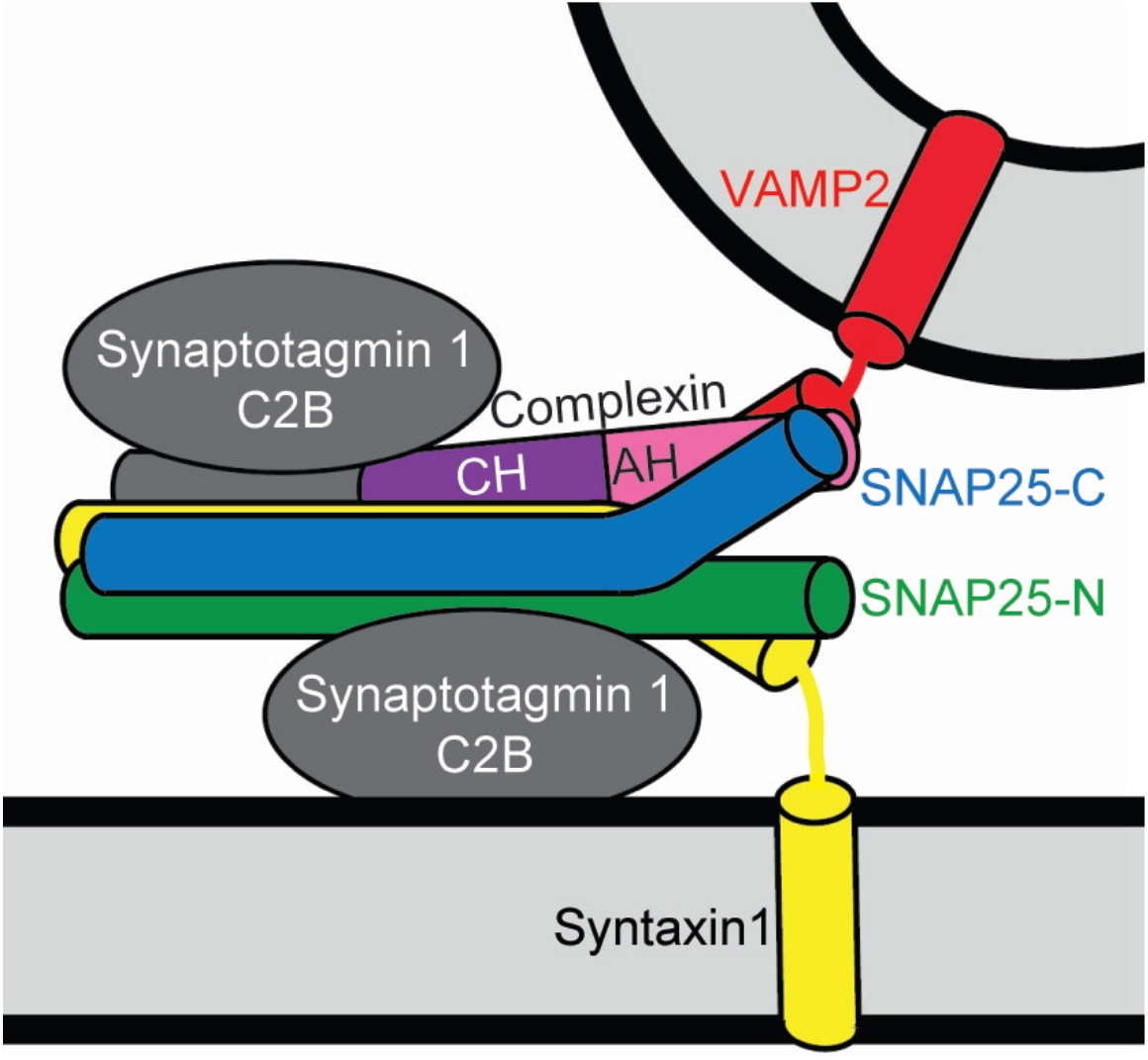
Model of how the Cpx accessory helix clamps fusion at the synapse. The central helix (CH) of Cpx stabilizes a partially zippered SNARE complex and the accessory helix (AH) binds the membrane-proximal C-terminal ends of SNAP-25 and VAMP2, preventing further SNARE complex zippering/assembly/membrane fusion.

## Experimental Procedures

### Constructs

Full-length t-SNARE complex (syntaxin 1A 1-288, His_6_-SNAP25 1-206): The bicistronic expression plasmid (pTW34) encoding untagged full-length rat syntaxin 1A (1-288) and N-terminally His_6_-tagged mouse SNAP25 (1-206) was described previously (Parlati et al., 1999). The N-terminal extension including the His_6_-tag is highlighted in red.

syntaxin1A 1-288 sequence: EIRGFIDKIAENVEEVKRK HSAILASPNPDEKTKEELEELMSDIKKTANKVRSKLKSIEQSIEQEEGLNRSSADLRIRKTQHSTLSRKFVEVMSEYNATQSDYRERCKGRIQRQLEITGRTTTSEELEDMLESGNPAIFASGIIMDSSISKQALSEIETRHSEIIKLENSIRELHDMFMDMAMLVESQGEMIDRIEYNVEHAVDYVERAVSDIKKAVKYQSKARRKKIMIIICCVILGIIIASTIGGIFG

His_6_-SNAP25B 1-206 sequence:

MRGSHHHHHHGSMAEDADMRNELEEMQRRADQLADESLESTRRMLQLVEESKDAGIRTLVMLDEQGEQLERIEEGMDQINKDMKEAEKNLTDLGKFCGLCVCPCNKLKSSDAYKKAWGNNQDGVVASQPARVVDEREQMAISGGFIRRVTNDARENEMDENLEQVSGIIGNLRHMALDMGNEIDTQNRQIDRIMEKADSNKTRIDEANQRATKMLGSG t-SNARE complex with C-terminally truncated SNAP25 (1-200/1-194): pTW34 (Parlati et al., 1999) was used as template DNA to truncate the SNARE motif at layer +6 (1-194) or around layer +8 (1-200) by introducing stop codons at the corresponding positions.

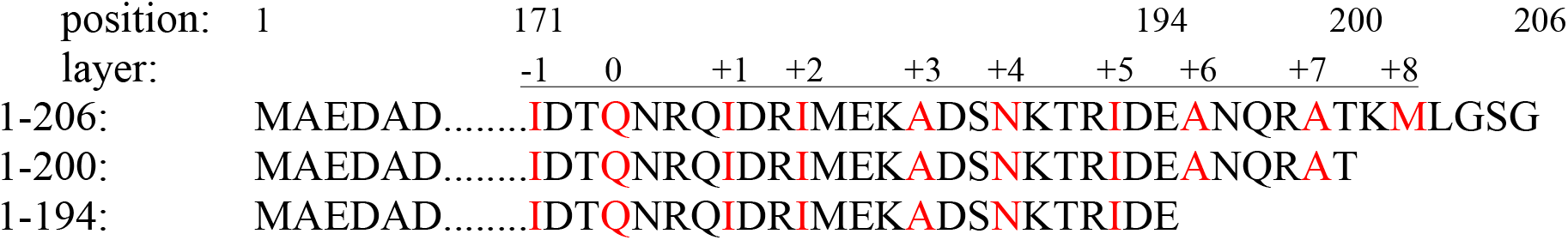

The amino acid positions and the hydrophobic layers (highlighted in red) of the SNAP25 SNARE motif are numbered and indicated above the sequences.

VAMP2 (1-116): A DNA construct encoding GST-tagged mouse VAMP2 (pSK28) was described previously (Kedar et al., 2015). The GST-tag was removed by thrombin cleavage resulting in a short N-terminal extension highlighted in red.

GSMSATAATVPPAAPAGEGGPPAPPPNLTSNRRLQQTQAQVDEVVDIMRVNVDKVLERDQKLSELDDRADALQAGASQFETSAAKLKRKYWWKNLKMMIILGVICAIILIIIIVY FST

synaptotagmin1: A DNA construct (pLM6) encoding His_6_-tagged rat Syt1 lacking the lumenal domain was described previously (Mahal et al., 2002). To replace the C-terminal His_6_-tag of this construct with a twin-Strep-tag (highlighted in red), the construct was subcloned into pPSG IBA103 resulting in pSK151.

MGPWALIAIAIVAVLLVVTSAFSVIKKLLFKKKNKKKGKEKGGKNAINMKDVKDLGKTMKDQALKDDDAETGLTDGEEKEEPKEEEKLGKLQYSLDYDFQNNQLLVGIIQAAELPALDMGGTSDPYVKVFLLPDKKKKFETKVHRKTLNPVFNEQFTFKVPYSELGGKTLVMAVYDFDRFSKHDIIGEFKVPMNTVDFGHVTEEWRDLQSAEKEEQEKLGDICFSLRYVPTAGKLTVVILEAKNLKKMDVGGLSDPYVKIHLMQNGKRLKKKKTTIKKNTLNPYYNESFSFEVPFEQIQKVQVVVTVLDYDKIGKNDAIGKVFVGYNSTGAELRHWSDMLANPRRPIAQWHTLQVEEEVDAMLAVKKGSAWSHPQFEKGGGSGGGSGGSAWSH PQFEK

Complexin II wt: A DNA construct (pMDL80) encoding His_6_-tagged human CpxII was described previously (Malsam et al., 2012). The His_6_-tag was removed by thrombin cleavage (short N-terminal extension highlighted in red).

GSHMDFVMKQALGGATKDMGKMLGGEEEKDPDAQKKEEERQEALRQQEEERKAKHARMEAEREKVRQQIRDKYGLKKKEEKEAEEKAALEQPCEGSLTRPKKAIPAGCGDEEEEEEESILDTVLKYLPGPLQDMFKK

Complexin II BPA mutants: For site-directed insertion of the unnatural amino acid p-benzoyl-phenylalanine (BPA), the corresponding codons of the CpxII sequence were replaced with the amber stop codon (TAG) using pMDL80 as template DNA.

Complexin II VAMP2- and SNAP25 binding mutants: pMDL80 was used as template DNA to generate the VAMP2- (K33E, R37E, A40K, Q44E, resulting in pSK130) and the SNAP25 (D27R, A30R, Q31E, E34R)-binding mutants.

### Protein expression and purification

All proteins were expressed in *E.coli* BL21(DE3). Cell disruption was performed with the high pressure pneumatic processor 110L (Microfluidics). The concentration of purified proteins was determined using Coomassie blue-stained SDS-PAGE employing BSA as standard protein and ImageJ software for quantification. Light chains of botulinum neurotoxins B and D expression vectors were kind gifts of Dr. Thomas Binz and late Dr. Heiner Niemann. t-SNARE expression and purification was performed as described previously (Parlati et al., 1999). VAMP2 was expressed and purified as described previously (Kedar et al., 2015). Synaptotagmin1 containing a C-terminal twin-strep-tag was expressed and purified as described previously for the His_6_-tagged protein (pLM6) (Malsam et al., 2012) with the following modifications: imidazole was omitted from all buffers and 50 mM biotin was used for protein elution from Strep-Tactin XT superflow high capacity resin. Complexin II wt and the Complexin II SNARE-binding mutants were expressed and purified as described previously (Malsam et al., 2012).

Complexin II BPA mutants were expressed using the pEVOL/pET system (Young et al., 2010). *E. coli* BL21(DE3) was co-transformed with the pEVOL-pBpF plasmid and the pET15b expression plasmid carrying the respective CpxII amber mutant. 1 liter LB media supplemented with a 50 mM potassium phosphate buffer (pH 7.3) was inoculated to grow a bacterial log-phase culture to an OD of 0.6. The culture was then transferred into a 2 liter beaker glass with a magnetic stir bar to rapidly chill to 30°C using an ice-water-bath and supplemented with 1 mM BPA by the addition of 10 ml 100x BPA stock solution (100 mM BPA dissolved in 100 mM NaOH). The culture was then transferred back into a 5 liter conical shake flask and incubated at 30°C. At an OD of 1.0 arabinose was added to a final concentration of 30 mM to induce tRNA transcription and expression of BPA-aminoacyl-tRNA synthetase for 1 hour at 30°C, followed by the induction of complexin expression with 0.3 mM IPTG for another 4 hours at 30°C. Purification of all BPA-containing complexin mutants was performed as described for Complexin II wt.

### Antibodies

The following antibodies were used: anti-CpxI/II rabbit polyclonal antibodies SM195 and # 122002 (Synaptic Systems), anti-VAMP2 rabbit polyclonal antibody #134 generated against amino acids 2-28 of VAMP2, anti-syntaxin1 mouse monoclonal antibody HPC-1, anti-Strep-tag II epitope mouse monoclonal antibody # 34850 (QIAGEN), anti-6xHis-tag mouse monoclonal antibody # H1029 (Sigma-Aldrich), anti-tubulin # T8660 (Sigma-Aldrich), Alexa Fluor 680 goat anti-rabbit # A21109 (Invitrogen), IRDye 800CW Goat anti-mouse # 926-32210 (LI-COR), horseradish peroxidase-conjugated goat antibodies directed against rabbit IgG # 111-035-003 and mouse IgG # 115-035-146 (Jackson ImmunoResearch Laboratories).

### Protein reconstitution into liposomes

Fluorophore-labeled lipids (Atto488-DPPE and Atto550-DPPE) were obtained from ATTO-TEC. ^3^H-DPPC (^3^H-1,2-dipalmitoyl-sn-glycero-3-phosphocholine) was from Amersham Pharmacia Biotech. All other lipids were from Avanti Polar Lipids. VAMP2/Syt1 lipid mix (3 μmol total lipid): 15 mol% DOPS (1,2-dioleoyl-sn-glycero-3-phosphoserine), 28.5 mol% POPC (1-palmitoyl-2-oleoyl-sn-glycero-3-phosphocholine), 25 mol% POPE (1-hexadecanoyl-2-octadecenoyl-sn-glycero-3-phosphoethanolamine), 5 mol% liver PI (L-α-phosphatidylinositol), 25 mol% cholesterol (from ovine wool), 0.5 mol% Atto488 1,2-dipalmitoyl-sn-glycero-3-phosphoethanolamine, 0.5 mol% Atto550 1,2-dipalmitoyl-sn-glycero-3-phosphoethanolamine) and trace amounts of ^3^H-DPPC (1,2-dipalmitoyl-sn-glycero-3-phosphocholine). Syntaxin1A/SNAP25 lipid mix (5 μmol total lipid): 15 mol% DOPS (1,2-dioleoyl-sn-glycero-3-phosphoserine), 35 mol% POPC (1-palmitoyl-2-oleoyl-sn-glycero-3-phosphocholine), 20 mol% POPE (1-hexadecanoyl-2-octadecenoyl-sn-glycero-3-phosphoethanolamine), 4 mol% liver PI (L-α-phosphatidylinositol), 1 mol% brain PI(4,5)P2 (L-α-phosphatidylinositol-4,5-bisphosphate), 25 mol% cholesterol (from ovine wool). Proteo-liposomes were generated by detergent dilution and purified as described previously (Kedar et al., 2015). Protein:lipid ratios: t-SNARE 1:1000, VAMP2 1:350, Syt1 1:700.

### Preparation of giant unilamellar t-SNARE vesicles

t-SNARE GUVs were generated from t-SNARE SUVs as described previously (Kedar et al., 2015).

### Fusion assays

Fusion reactions and data analysis were performed as described previously (Malsam et al., 2012).

### Liposome cross-linking

All solutions were adjusted to 280 mOsm. t-SNARE GUVs (250 μM lipid) were mixed with VAMP2-Syt1 SUVs (50 μM lipid) and 6 μM CpxII wt or BPA mutants in fusion buffer (20 mM MOPS-KOH, pH 7.4, 135 mM KCl, 0.5 mM MgCl_2_, 100 μM EGTA-KOH, 5 mM glutathione, 1 mM DTT) in a final volume of 100 μl and incubated for 30 minutes on ice. To remove unbound SUVs and complexin, the reaction-mix was overlayed on 100 μl of 70 mM sucrose in fusion buffer and underlayed with a cushion of 5 μl 250 mM sucrose in 1 mM HEPES-KOH, pH 7.4. GUVs were re-isolated by centrifugation at 10.000 x g for 15 minutes in a swinging-bucket rotor. The supernatant was removed and sedimented liposomes were gently resuspended in a total volume of 15 μl for subsequent cross-linking. Samples were irradiated on ice with a UV-LED lamp (Opsytec Dr. Gröbel GmbH) at 365 nm applying 15 pulses of 1 second at maximum irradiation power (25 W/cm²), inserting pauses of 2 seconds between pulses.

### Protease treatment of liposomes with Botulinum neurotoxins

Liposome samples were prepared and Cpx R37-BPA arrested SUV-GUV complexes re-isolated as described above for liposome cross-linking. 15 μl sedimented liposomes were diluted with 20 μl fusion buffer and heated at 98°C for 1 minute, followed by the addition of 10 μl 2% (w/v) Triton X-100. Solubilized proteins were subjected to site-specific proteolytic cleavage of VAMP2 with 0.5 μM BoNT/B or BoNT/D in final volumes of 50 μl at 37°C for 15 minutes. Samples were then mixed with 500 μl cold acetone and incubated for 3 hours at −20°C. Precipitated proteins were sedimented at 0°C for 30 minutes at 20.000 x g. Pellets were air-dried, solubilized in SDS-PAGE sample buffer and proteins separated on 16% Tris-Tricine gels for Coomassie brilliant blue staining or Western-blot analysis.

### Cryo-electron microscopy and cryo-tomography

Samples were processed for plunge freezing as described in (Malsam et al., 2012). Samples were imaged at cryo temperatures under standard low-dose conditions on either a FEI Tecnai Spirit electron microscope (120 kV, 23000x) equipped with a Gatan Ultrascan 4000 CCD camera (4.9 Å/pixel, Gatan, Pleasanton CA) or on a FEI Tecnai Polara electron microscope (300 kV, 23000x) equipped with a Falcon 2 direct electron detection camera (4.85 Å/pixel). Imaging of each sample was performed in an unbiased, semiautomated manner using SerialEM (Bharat et al., 2014; Mastronarde, 2005). The number of SUVs bound to one GUV was recorded for each sample.

For morphological analysis of prefusion sites, cryo-tomograms were automatically collected on a FEI Titan Krios electron microscope operated at 300 kV and equipped with a Gatan Quantum 967 LS energy filter and a K2 xp direct electron detector. Dose-symmetric tilt series (Hagen et al., 2017) were acquired in EFTEM mode with a 20eV slit and a calibrated magnification of 130kx. A tilt range of ±66 with 3° angular increments was chosen, resulting in a total dose of approximately 130 electrons/Å^2^. Super-resolution frames were aligned on-the-fly with a frame-alignment algorithm built into SerialEM (Mastronarde, 2005) and Fourier cropped to 4k × 4k images giving a pixel size of 1.05 Å/pixel. Subsequently, tilt series were sorted and filtered by cumulative electron dose (Grant and Grigorieff, 2015; Schur et al., 2016). Tomograms were generated by weighted-back projection using the IMOD/etomo pipeline (Kremer et al., 1996). For visual inspection, tomograms were binned twice and low-pass filtered resulting in a final pixel size of 4.2 Å/pixel. Medium magnification grid square maps (6500x, 2.27 nm/pixel) generated during the imaging sessions were also used for docking analysis.

### Preparation of cultured hippocampal neurons

Murine microisland cultures were prepared as described (Xue et al., 2007). CpxI-III triple KO neurons were described previously (Xue et al., 2008). Animals were handled according to the rules of Berlin authorities and the animal welfare committee of the Charité – Universitätsmedizin Berlin, Germany. Primary hippocampal neurons were prepared from mice on embryonic day E18 and plated at 300 cm^−2^ density on WT astrocyte microisland for autaptic neuron electrophysiology. For Western blotting hippocampal neurons were plated at 10.000 cm^−2^ on continental WT astrocyte feeder layer.

### Lentiviral constructs and virus production

For expression of CpxII variants within neuronal cells, a modified lentiviral vector (Lois et al., 2002) was used containing a human Synapsin-1 promoter, driving the expression of a nuclear GFP and the CpxII variant (WT CpxII, VAMP2- or SNAP25-binding mutants of CpxII). The cDNAs were coupled via a self cleaving P2A site (Kim et al., 2011) leading to bicistronic expression of the 2 proteins (f(syn)NLS-GFP-P2A-CpxII-WPRE). Lentiviral particles were prepared by the Charité Viral Core Facility as previously described (Lois et al., 2002), vcf.charite.de. Briefly, HEK293T cells were cotransfected with the shuttle vector f(syn)NLS-GFP-P2A-CpxII-WPRE and helper plasmids, pCMVdR8.9 and pVSV.G with polyethylenimine. Virus containing supernatant was collected after 72 h, filtered, aliquoted, flash-frozen with liquid nitrogen, and stored at −80°C. For infection, about 5×10^5^–1×10^6^ infectious virus units were pipetted onto 1 DIV hippocampal CpxI-III triple KO neurons per 35 mm-diameter well.

### Electrophysiology of cultured neurons

Whole cell patch-clamp recordings in autaptic glutamatergic neurons were performed as previously described(Trimbuch et al., 2014). The extracellular solution contained 140 mM NaCl, 2.4 mM KCl, 10 mM Hepes, 2 mM CaCl_2_, 4 MgCl_2_, 10 mM Glucose (pH adjusted to 7.3 with NaOH, 300 mOsm). The patch pipette solution contained 136 mM KCl, 17.8 mM Hepes, 1 mM EGTA, 0.6 mM MgCl_2_, 4 mM ATP-Mg, 0.3 mM GTP-Na, 12 mM phosphocreatine and 50 units/ml phosphocreatine kinase (300 mOsm, pH 7.4). Neurons were clamped at −70 mV with a Multiclamp 700B amplifier (Molecular Devices) under control of Clampex 9 (Molecular Devices) at DIV 14-17. Data were analyzed offline using Axograph X (AxoGraph Scientific) and Prism 7 (GraphPad Software). Statistic significances were determined by one-way analysis of variance (ANOVA) with Kruskal-Wallis test followed by Dunn’s post test to compare all groups.

EPSCs were evoked by a brief 2 ms somatic depolarization to 0 mV. EPSC amplitude was determined as the average of 6 EPSCs at 0.1 Hz. RRP size was determined by measuring the charge transfer of the transient synaptic current induced by a pulsed 5 s application of hypertonic solution (500 mM sucrose in extracellular solution). Pvr was calculated as the ratio of the charge from an evoked EPSC and the RRP size of the same neuron. Evoking 5 or 50 synaptic responses at 50 or 10 Hz respectively in standard external solution analyzed short-term plasticity. For analyzing mEPSCs, traces were digitally filtered at 1 kHz offline. Then the last 9 seconds of 5 traces of EPSCs at 0.1 Hz were analyzed using the template-based mEPSC detection algorithm implemented in Axograph X (AxoGraph Scientific) and substracted from background noise by detecting events in the last 4 seconds of 5 EPSCs at 0.2 Hz in 3 mM NBQX (Tocris) in extracellular solution.

## Acknowledgments

We thank Peter Schultz for providing us the pEvol system to introduce the unnatural amino acids into proteins and Timon Braun and Susanne Kreye for technical assistance. This work was funded by the Deutsche Forschungsgemeinschaft (DFG; German Research Foundation) - project number 112927078 - TRR83 to J.M., A.S., I.S., and T.H.S., project number – 278001972 - TRR186 to S.B., J.M., A.S., T.T., F.Z, T.H.S., and C.R.) and the European Research Council (ERC) under the European Union’s Horizon 2020 research and innovation programme ERC-CoG-648432 MEMBRANEFUSION (to J.A.G.B).

## Author contributions

J.M., C.R., I.S., J.A.G.B. and T.H.S. designed the study. J.M, S.B, A.S. performed the biochemical and proteo-liposomes experiments. T.T. and F.Z. conducted the electrophysiological experiments. A.F.-P.S. performed the cryo-electron microscopy and cryo-tomography experiments. K.W. provided critical insights to conceive the structural model and to select appropriate mutants to prove the model. All authors performed data analysis, discussed the results, provided critical feedback on the experiments, and edited the final manuscript.

