## Supplemental information for "Complexin suppresses spontaneous exocytosis by capturing the membrane-proximal regions of VAMP2 and SNAP25"

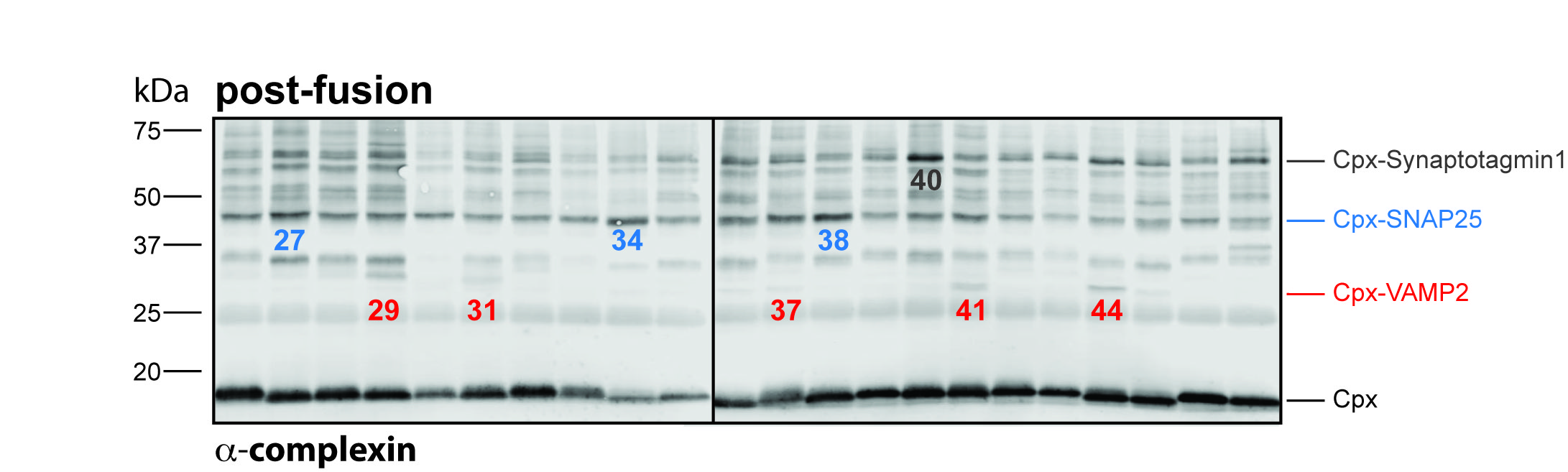

Figure S1. Related to Figure 2; Postfusion CpxII cross-link products. GUVs containing the full-length t-SNARE complex (syntaxin1/SNAP25) were mixed with SUVs containing synaptotagmin1 (Syt1) and the v-SNARE VAMP2 in the presence of Cpx wt or Cpx-BPA mutants. A pre-incubation for 30 minutes on ice allows docking of SUVs to the GUVs by Syt1 and trans-SNARE complexes. Subsequently, membrane fusion was triggered by 100 µM Ca^2+^, followed by the irradiation of the samples at 365 nm for 15 seconds on ice. Cross-link products were identified by Western blot analysis using an anti-Cpx antibody. Identified crosslink products and their positions are indicated. Experiments were repeated two times, yielding virtually identical results.

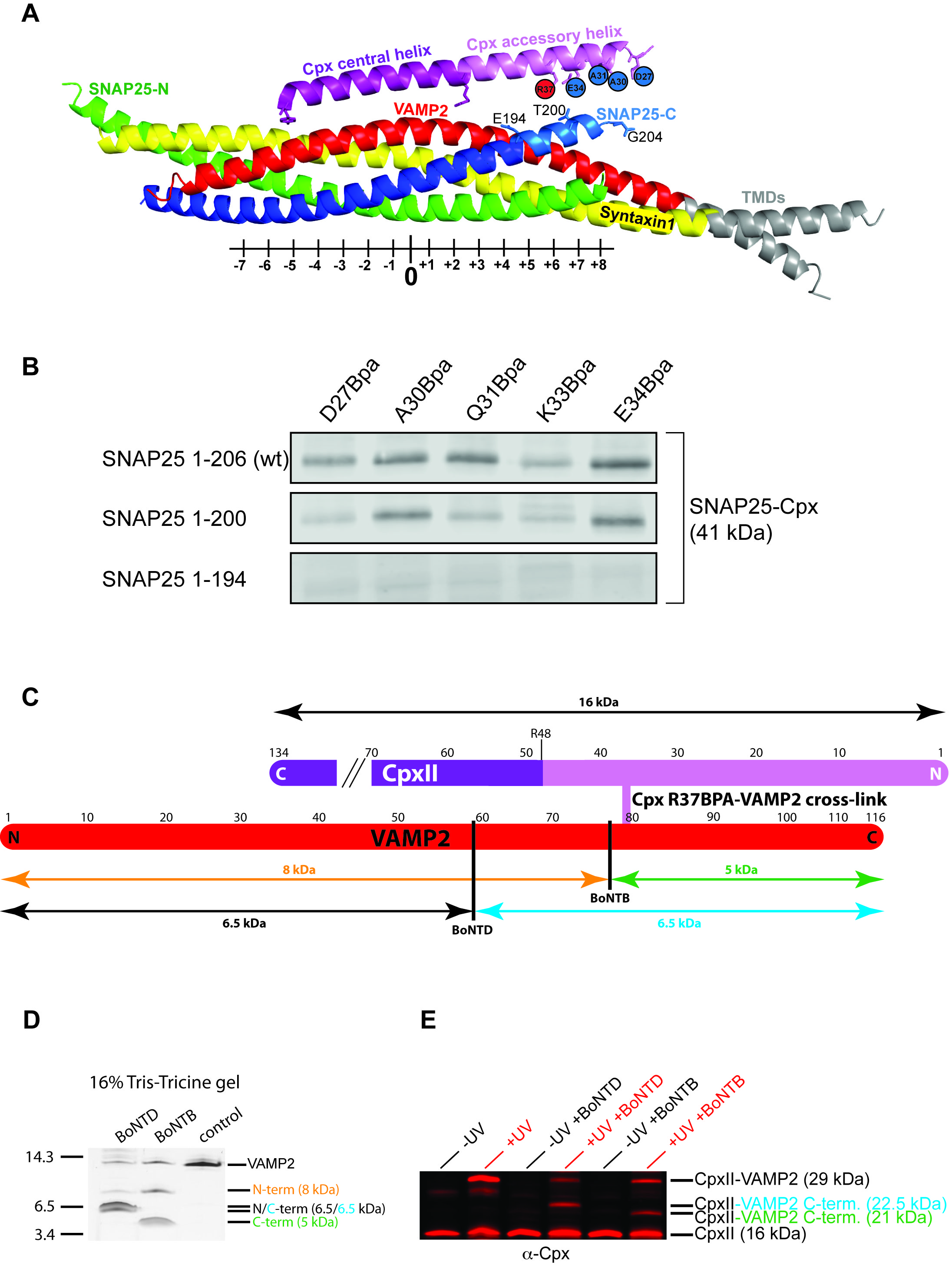

Figure S2. Mapping the Cpx accessory helix-SNARE binding region by truncation and proteolytic fragment analysis. SNAREpins containing Cpx BPA mutants at the indicated positions and t-SNARE complexes with either full-length (1-206) or two C-terminally truncated SNAP25 proteins were accumulated and subjected to UV-induced cross-link formation as described above. (A) depicts the employed constructs in a model of the postfusion structure (modified from Chen et al., 2002; Stein et al., 2009)). Highlighted in light blue is the C-terminal end of the SNAP25 helix. The last amino acids of the individual truncation constructs are labeled as E194 and T200, respectively. (B) CpxII-SNAP25 crosslinks still form with the SNAP25 truncation lacking the last six amino acids but are lost when using the 12 amino acid truncation as revealed by a complexin-specific antibody (n = 2). (C) To assign the Cpx accessory helix binding region to the N- or C-terminus of VAMP2-, SNAREpins containing Cpx R37-BPA were cross-linked and subjected to protease digestion using the VAMP2-specific Botulinum neurotoxins B and D, respectively. (C) and (D) As depicted BoNTB generates VAMP2 fragments of 8 and 5 kDa, respectively, while BoNTD produces two fragments with near identical length as confirmed by SDS-PAGE analysis and Coomassie blue staining. (E) To analyze UV- and protease-treated liposome samples for Cpx-VAMP2-fragment size, samples were processed for Western-blotting analysis and products were detected with a Cpx-specific antibody. A unique Cpx-VAMP2 fragment of 21 kDa is generated in a UV- and BoNTB treated sample consisting of the combined masses of Cpx (16 kDa) and the C-terminal 5 kDa fragment of VAMP2 (last lane). The data position the CpxII crosslinks to the C-terminal ends of VAMP2 and SNAP25.

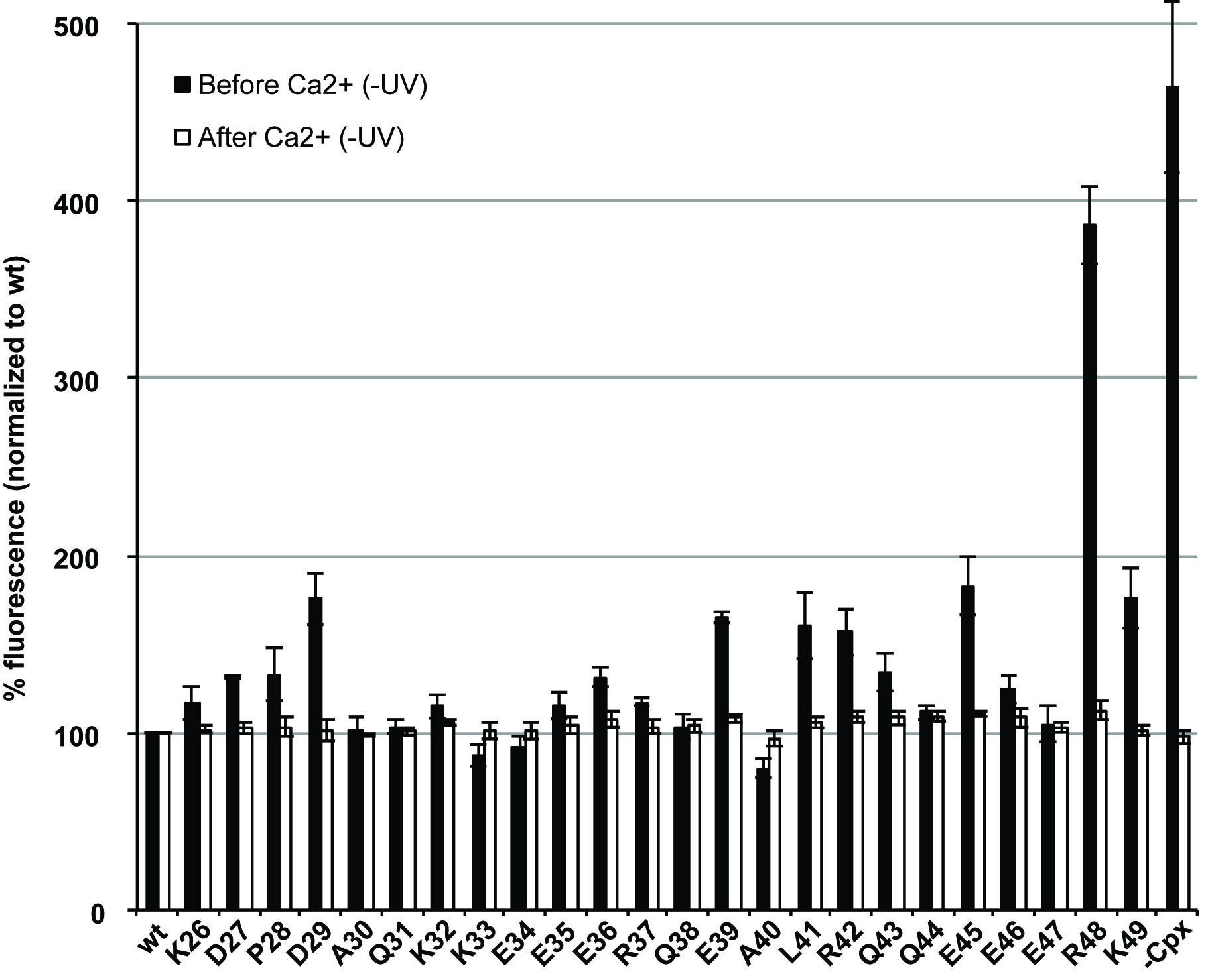

Figure S3. Related to Figure 3; Effect of Cpx-BPA mutants on membrane fusion. GUVs containing full-length t-SNARE complexes (syntaxin1/SNAP25) were mixed with SUVs containing Syt1 and the v-SNARE VAMP2 in the absence/presence of Cpx wt or Cpx-BPA mutants as indicated. A pre-incubation for 30 minutes on ice allows the docking of SUVs to GUVs by Syt1 and trans-SNARE complexes. Subsequently, fusion kinetics were recorded at 37°C for 1 minute in the absence of Ca^2+^. The measurements were continued for another two minutes after injection of 100 µM free Ca^2+^ to trigger fusion. Data were normalized to the fluorescence value of a control reaction containing complexin wt (before Ca^2+^, black bars) or to the maximum total fluorescence value after detergent lysis (after Ca^2+^, white bars). Please note that R48BPA impairs the binding of CpxII to the SNAREpins, abolishing any CpxII effects (similar to -Cpx). Error bars indicate s.e.m. (n = 3).

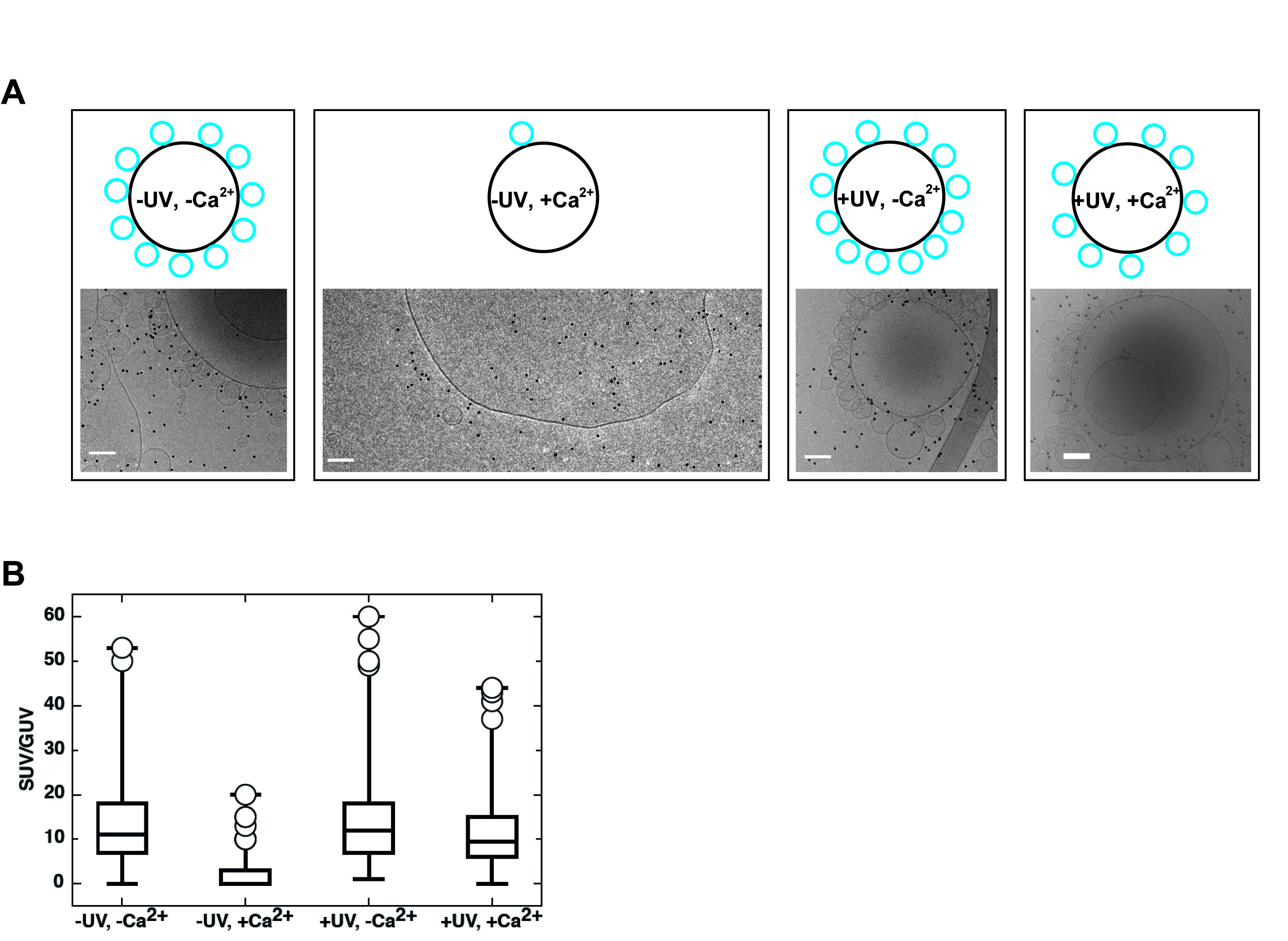

Figure S4. Related to Figure 3; Morphology of UV-arrested, calcium-resistant SUV-GUV docking sites. (A) VAMP2-Syt1-containing SUVs were allowed to dock to t-SNARE GUVs in the presence of Cpx-A40BPA and subjected to UV irradiation (control samples without UV treatment). Subsequently, fusion was triggered with 100 µM Ca^2+^ for 2 min at 37°C (control reactions without Ca^2+^) and samples were plunge-frozen for analysis by cryo-EM. Scale bar = 100 nm. (B) Box plot diagrams showing the average number of SUVs docked to one GUV. Box (with median) comprises 25%-75% of data, far outliers are drawn as circles, minimum and maximum values are drawn as whiskers. n = 348 (-UV, -Ca^2+^), 179 (-UV, + Ca^2+^), 407 (+UV, - Ca^2+^), 330 (+UV, + Ca^2+^).

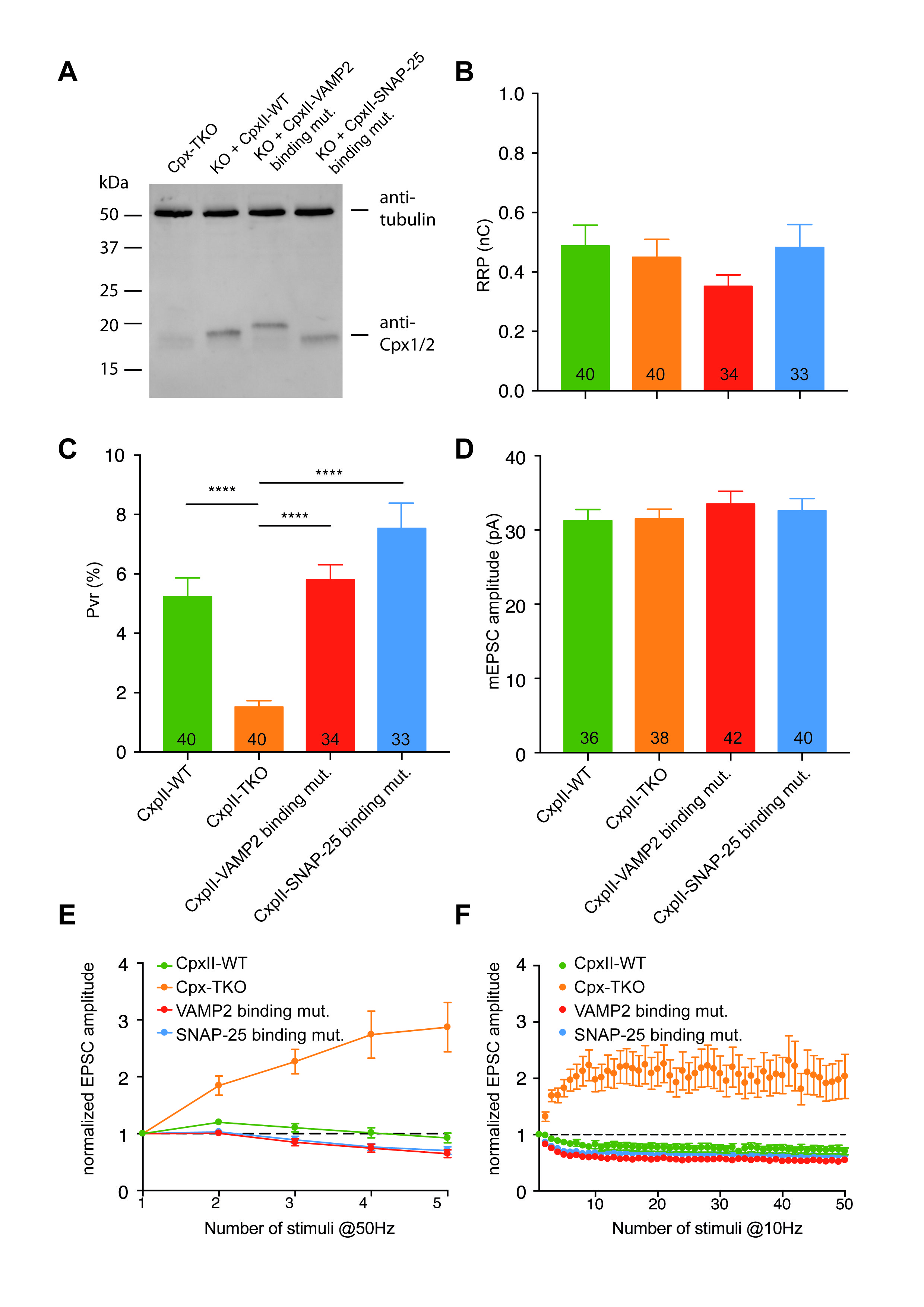

Figure S5. Related to Figure 4; Analyses of Cpx mutants affecting VAMP2 or SNAP25 binding in neurons. (A) Cpx triple knockout neurons were infected with lentivirus expressing wt or mutant complexins. Cells were processed for Western-blot analysis at DIV14 using the indicated antibodies. (B-D) Bar graph analysis of the readily-releasable pool size (RRP), the release probability (Pvr) and the mEPSC amplitude. The data were obtained from two independent cultures. The numbers in the bars indicate the sample size. Error bars indicate s.e.m. **** p<0.0001. (E and F) Analyzes of short-term plasticity after evoking 5 or 50 responses at 50 or 10 Hz respectively. Error bars indicate s.e.m. The n values for 50 Hz are: n(CpxII wt) = 41, n(CPX-TKO) = 41 , n(Vamp2 binding mut.) = 38, n(SNAP-25 binding mut.) = 40. The n values for 10 hz are: n(CpxII wt) = 41, n(CPX-TKO)=40 , n(Vamp2 binding mut.) = 38, n(SNAP-25 binding mut.) = 40.
